# Tailpiece Heating device for effective prevention of biofilm growth in sink plumbing

**DOI:** 10.1101/2023.09.15.558019

**Authors:** Shireen M Kotay, William Guilford, Brian A Pettygrove, Alan J Komisarcik, Samantha A. Hughes, Amy J Mathers

## Abstract

In recent years, numerous hospitals have linked patient infections to *Klebsiella pneumoniae* producing Enterobacterales (KPCE) and other resistant bacterial species in their wastewater systems and handwashing sinks. Wastewater plumbing provides a reservoir for bacteria, making them incredibly difficult to eliminate through traditional disinfection methods. Data suggests that patients become infected when bacteria grow or migrate up the proximal wastewater plumbing and into the sink basin, and are subsequently dispersed onto surrounding surfaces. Therefore, a novel electronic device was developed that acts at the highest risk area, just below the sink drain, to heat and dry out the biofilm and creating a biofilm barrier and prevent upward growth from the sink trap. The efficacy of the first prototype of a tailpiece heater (TPH) in preventing drain colonization was tested using GFP-expressing *Escherichia coli* (GFP-*E. coli*) as the challenge organism. In control sinks without the TPH, GFP-*E. coli* biofilm grew from the p-trap upwards to colonize the drain within 7 days. Sinks with a TPH set to 75°C were found to prevent sink drain colonization. In contrast, 65°C was not adequate to prevent drain colonization. Using KPCE the TPH was more effective than no heat control in preventing drain colonization in sinks over time. Lastly, when challenged with seeding from above, the TPH also effectively prevented KPCE colonization at the drain level. Heating of the tailpiece may offer a safe, effective, and economically attractive approach to preventing the spread of resistant bacterial species from contaminated drain biofilm to patients.

## INTRODUCTION

Hospital wastewater discharge systems have long been recognized to harbor pathogens and opportunistic bacteria. There is growing consensus that sink drains and the associated plumbing in the healthcare environment can act as reservoirs of pathogenic and multidrug resistant organisms, including Carbapenem-resistant Enterobacteriaceae which may spread to patients and surrounding surfaces (1, 2). With sinks as established sources of outbreaks, interventions to reduce the transmission of gram-negative bacilli from the environment to patients have been prioritized in an effort to protect patients (1, 3).

Interventions such as replacement of plumbing (4-9) and/or the use of chemical disinfectants, such as bleach (6), acetic acid (7), and hydrogen peroxide (5), have been previously attempted to contain sink-related transmissions. Despite limited success using these interventions the target organisms are difficult to eradicate, and their recurrence in the sinks following the intervention has often been reported (2, 10). This recurrence is associated with resilient biofilms formed on the luminal surface of sink drainage plumbing that may regrow quickly under favorable conditions. Self-disinfecting sinks that heat sink p-traps and modified sink construction to reduce splashing are new and emerging technologies, (11-13) but their effectiveness in real-world setting is yet to be fully understood.

Sink traps, also known as p-traps, form a water barrier against noxious sewer gas escaping to indoor air (Fig 1). The retained water in the p-traps provides favorable conditions for microorganisms to survive and promotes their propagation as resilient biofilms. Further, the p-trap and its connections to the hospital wastewater system may experience some bidirectional flow (14). It can therefore potentially be inoculated from two directions: i) from above due to organisms disposed into the sink drain via patients’ bodily fluids etc. or ii) due to retrograde back-flow from the sanitary stack and connecting pipes (Fig 1). These mechanisms for establishing multidrug resistant gram-negative bacteria in the sinks has been experimentally demonstrated previously (14-16). From a microbiology standpoint the sink wastewater system can be divided into three zones. First, the ‘Inlet, or Transmission zone’, that comprises the sink bowl, faucet, drain and the upper portion of the tailpiece and is subject to inoculation from above. Further, it is from this zone that transmission to patients could happen if the drain and/or sink bowl is colonized (Fig 1). Second, the ‘Reservoir zone’ is comprised mainly of the p-trap, which is the holding point for the water, lower portion of the tailpiece and proximal portion of the trap arm. This zone is a potential breeding ground for drug-resistant pathogens, providing organisms both with favorable growth conditions and potential selective pressures from antimicrobials and disinfection soaps (e.g. chlorhexidine). And finally, the ‘Outlet, or Intermixing zone’, is the trap-arm that carries outflow from the p-trap to the connected wastewater plumbing downstream. This zone offers an opportunity for intermixing and backflow and both inoculates and is inoculated by the distal wastewater system.

**Figure 1:**
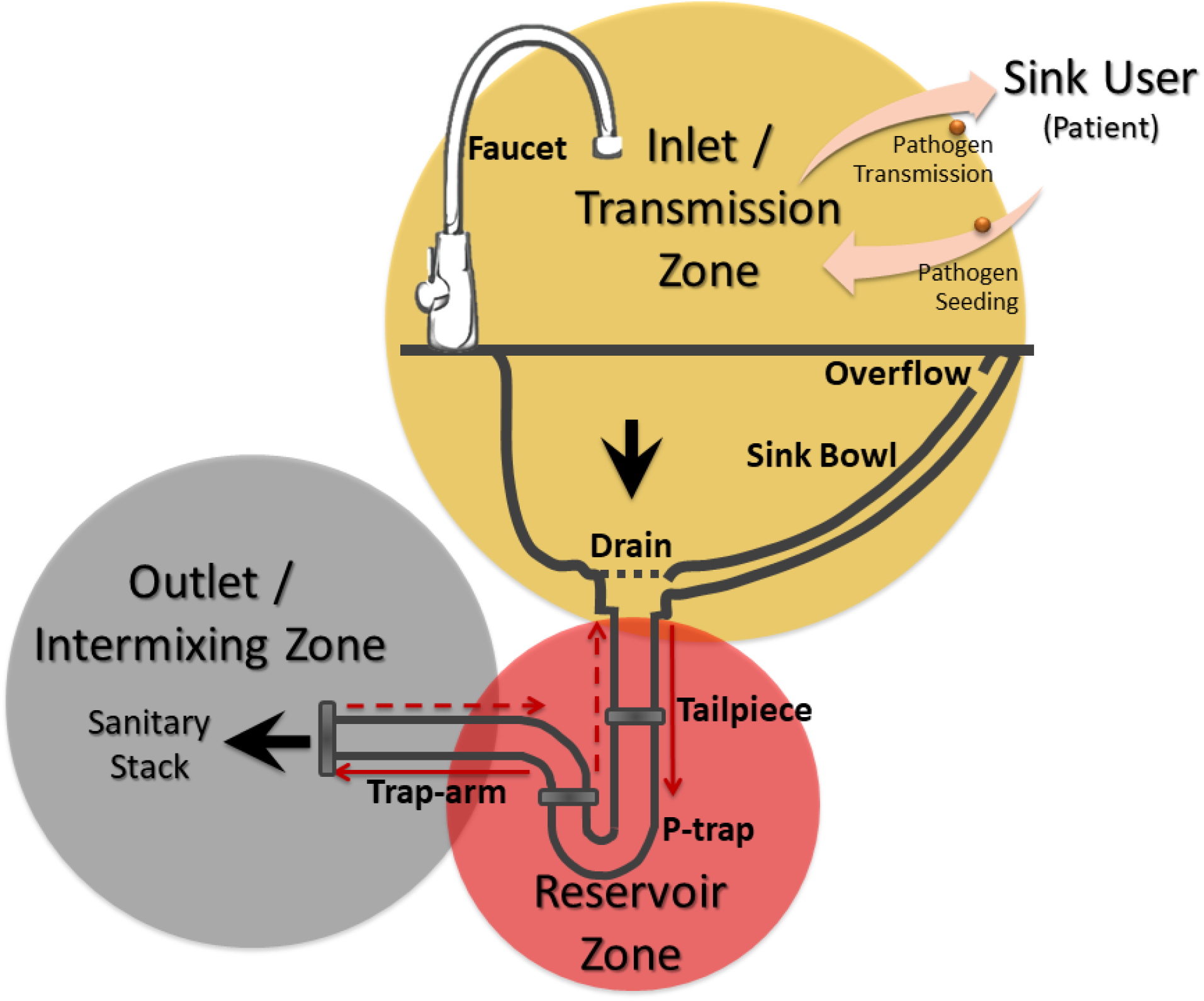
Conceptual infographic depicting the mechanism of sink microbial seeding, colonization, intermixing via wastewater plumbing and transmission.

Wastewater systems are designed such that all waste flows down from the sink or toilet, however, moist environments favor establishment and spread of biofilm inside the plumbing even against gravity (14, 16). With this knowledge and an aim to mitigate transmission from sink drain biofilms to the patient care environment, a scientifically informed electronic device was developed. We employ a strategy to target the highest risk area, biofilm just below the sink drain, with an objective to use heat to dry the biofilm and create a biologic barrier to prevent the upward growth from the sink trap to the drain. In this study, we challenge the device in an *ex situ* controlled environment to determine its efficacy in preventing drain colonization.

## MATERIALS AND METHODS

### Sinks and settings

A dedicated sink gallery was used for the testing of the first prototype of a tailpiece heater (TPH) targeting the biofilm right below the drain. Design and operation of these sinks in this *ex situ* setting was described previously (14, 15). Testing of subsequent prototypes of the TPH was performed in a Biosafety Level 2 certified sink lab with modular sinks. Design and operation of these sinks was described in the earlier study (17). For the operation, sinks were automated such that water (1.3L) at approximately 5 L/min and soap (1.5ml) flowed for 15 seconds, once every two hours.

### Tailpiece heater design and construction

Assembly of the prototype TPH is shown in Fig 2. The prototype device was built around a 1.25 inch (32mm) 20 gauge chromed brass tailpiece (Dearborn Brass^®^-Oatey, Cleveland, Ohio). Self-adhesive resistive heat tape (Clayborn Lab, Truckee, CA; 12 volt, 33 watt) was wrapped around the tailpiece in a helical fashion to cover a length of approximately 75 mm. An electronic temperature sensor (TMP36, Analog Devices) and an irreversible thermal cutoff fuse (Selco SWTC-262-3535, Allied Electronics; 128°C fault) were attached to the outer surface of the tailpiece in the middle of the heated region using Kapton tape. The thermal cutoff was wired in series with the heat tape as a failsafe against runaway heating. The heater assembly was surrounded by an insulating layer of fire-retardant silicone foam (McMaster-Carr). A microcontroller project board (Arduino Pro Micro, Sparkfun) controlled a MOSFET (Fairchild FQP30N06L) to control current through the heating element. Temperature was regulated through a software Proportional – Integral – Derivative (PID) controller implemented on the microcontroller, and the status of the thermal cutoff fuse was continuously monitored by the voltage drop across the heating element. In addition to the thermal cutoff fuse, as an added safety measure the microcontroller triggered a reversible cooldown to room temperature if the tailpiece temperature were to rise above 85° C.

**Figure 2.**
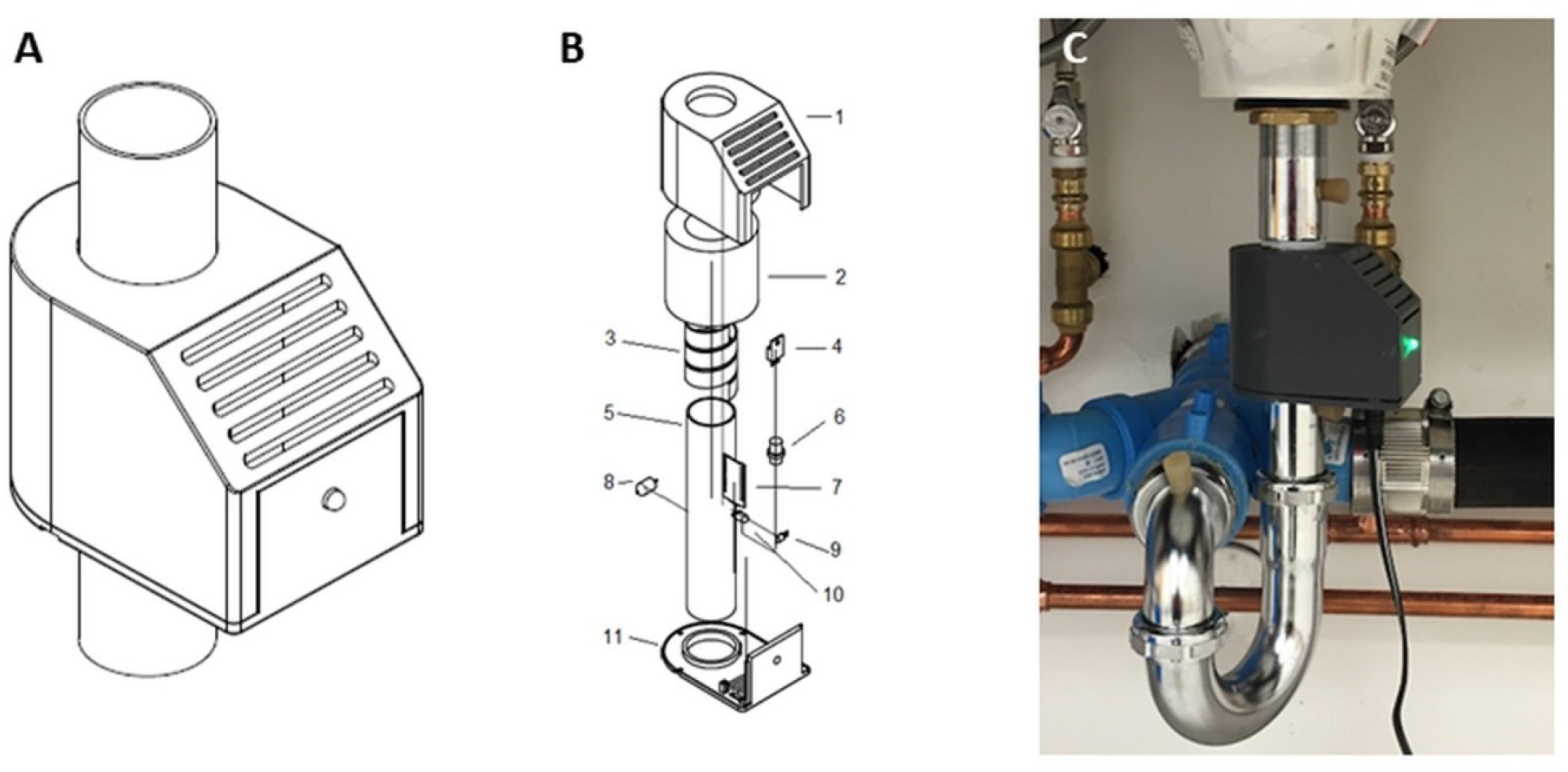
**(A)** Assembled prototype tailpiece heater on a 1.25 inch sink tailpiece. (**B**) Exploded view of the TPH. 1) upper case; 2) silicone foam insulation; 3) heating element; 4) MOSFET; 5) tailpiece; 6) power connector; 7) microcontroller circuit board; 8) thermal cutoff fuse; 9) temperature sensor; 10) LED; 11) lower case. **(C)** Prototype I installed on sink.

The entire assembly was surrounded by a protective case 3D printed from ABS plastic. The USB port of the microcontroller was accessible without disassembling the device. This allowed for monitoring and modification of the device’s firmware. Vents were included on the top and bottom surfaces of the electronics chamber to keep the electronics cool. Also included on the case was a tri-color LED to indicate operational and error status.

### Tailpiece heater installation and settings

For uniformity and comparison, tailpieces of same diameter and length were installed in sinks with and without TPH devices (Fig 3). TPH devices were assembled 40 mm from the end of the tailpiece – approximately 100 mm from the upper surface of the drain once installed. Tailpieces with and without heaters were fitted with sampling ports and rubber stoppers 25 mm above and below the heater to accommodate sampling the luminal surfaces (Fig 3 B, C). Each sink basin and counter were surface sterilized prior to start of the experiment. New strainers (drains) were also installed prior to each experiment (Fig 3A). Heat settings and cycle were controlled via the firmware. Two temperature settings on the TPH, 65°C and 75°C were tested in this testing. In each case the cycle consisted of 1 hour of heating at the target temperature, followed by 3 hours of no heat allowing cooling back to the room temperature.

**Figure 3.**
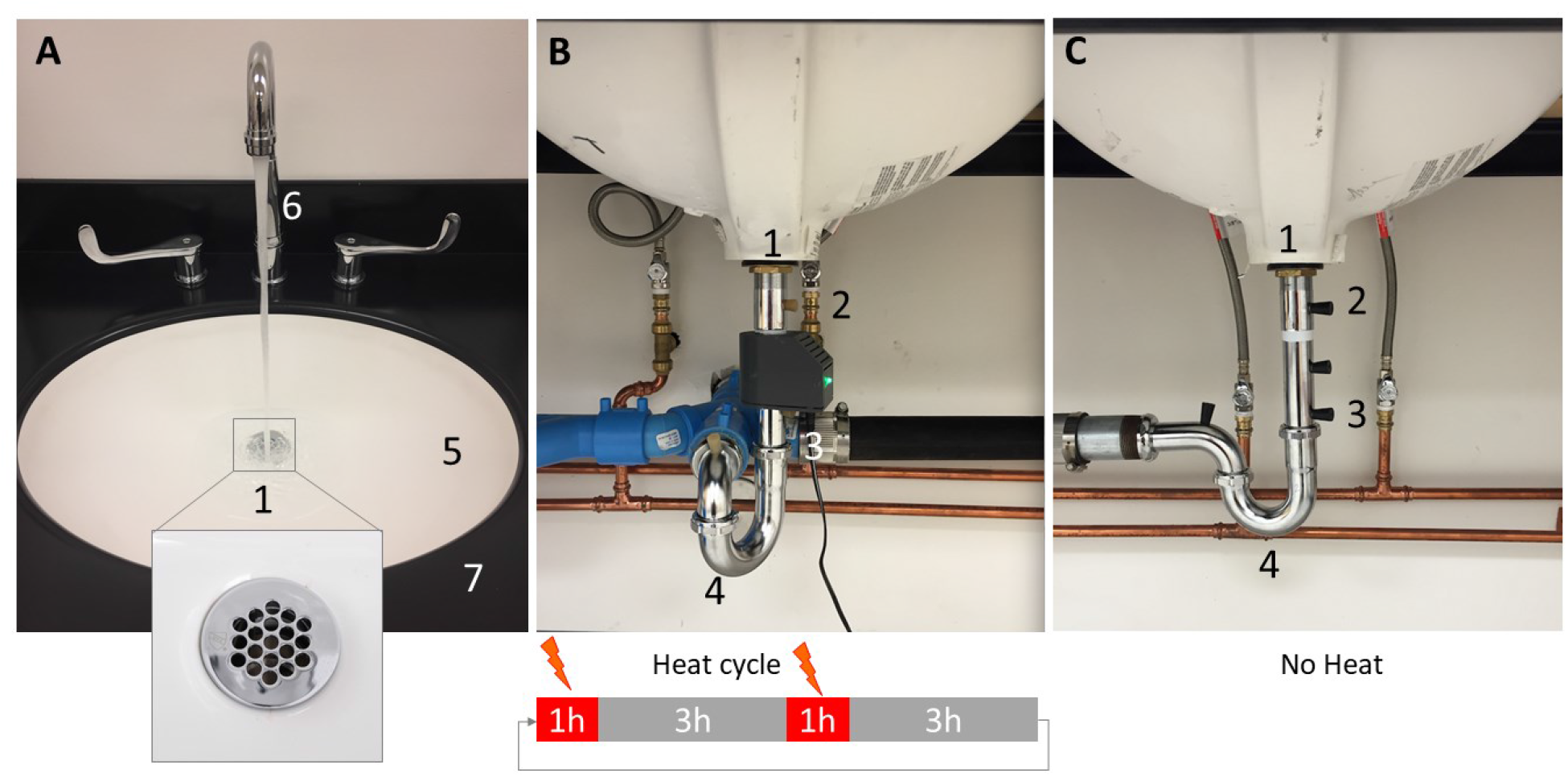
**(A)** Handwash sink top view - drain plate enlarged in the inset **(B)** Sink with the tailpiece heater, **(C)** Sink without tailpiece heater. Parts of the sink and sampling locations highlighted as 1-Drain, 2-Sampling port on tailpiece above heat, 3-Sampling port on tailpiece below heat, 4-P-trap, 5 – Sink bowl, 6-Faucet, and 7-sink counter. Sampling ports on the tailpiece were custom-drilled to provide access to the luminal surfaces of the fixtures. The heating cycle was 1 hour-ON, 3 hours -OFF.

### Inoculation, sampling and enumeration of challenge organisms

For the efficacy testing of the first prototype of the TPH, GFP-expressing *Escherichia coli* was used as the challenge organism. For the testing of subsequent prototypes, *Klebsiella pneumoniae* producing Enterobacterales (KPCE) isolates previously isolated from the hospital sink drain or P-trap were used as the challenge organisms. Methods as described in previous studies (14, 15, 17) were followed for growth and inoculation of the bacterium into sink plumbing fixtures.

For inoculation in P-trap, 10ml mid-log phase culture of the respective challenge organism (10^9^ CFU/ml) was added into the P-trap water (∼150ml) through the lower-most sampling port on the tailpiece using a 60ml syringe attached to silicone tubing (Cole-Parmer, Vernon Hills, IL). The inoculum was mixed with the P-trap water by repeated withdrawal and injection of the inoculum, with precautions taken to avoid unintentional inoculation of drain (strainer) or sink bowl (basin). For the drain inoculation (seeding from above), a 10ml mid-log phase culture of the respective challenge organism (10^9^ CFU/ml) was evenly applied on the surface of the drain plate using a sterile pipet. Following inoculation, 25ml tryptic soy broth (TSB; BD™ Bacto™) followed by 25ml (x2) 0.85% saline was added on a daily basis through the drain to promote growth.

To monitor the growth of the challenge organisms inside the plumbing, sterile cotton swabs (Covidien™, Mansfield, MA) presoaked with 0.85% sterile saline were inserted through the sampling ports and swab were turned in a circular motion on the luminal surface (∼20 cm^2^). Sample swabs were pulse-vortexed in 3ml saline and serial dilutions were plated. In case of GFP-*E*.*coli* experiments samples were plated on Tryptic Soy Agar (TSA) and for KPCE experiments samples were plated on Colorex™ mSuperCARBA™ agar (Northeast Laboratories, Waterville, ME). Colored and non-colored colonies with unique morphology were further processed for species identification via the VITEK2 (Biomerieux, Durham, NC) as described previously (11). A single colony isolated was considered positive for GFP-*E. coli* or KPCE and formal quantitation was not performed.

## RESULTS

### TPH Device Safety testing

Two tests were conducted to validate the efficacy of the controller. First we demonstrated the PID software controller’s ability to reach and maintain the initial target temperature of 75° C for sixty minutes with very little fluctuation (Figure 4A). For the first 40 seconds of a heating cycle, the tailpiece rose in temperature at rate of 0.7 °C/sec (Figure 4B); a linear relationship is expected under PID control when the temperature is far from its target. After that the temperature rose exponentially toward its target with a time constant of 64 seconds. The typical time to reach 75 °C was approximately 4.5 minutes. Second, we tested the device’s response to cold water flowing through the tailpiece – a circumstance that would inevitably be experienced in a clinical setting. The PID controller responded adequately to the temperature perturbations caused by the flow of cold water at each phase (Figure 4C). In cases where the heating cycle was interrupted by cold water flow, an equivalent amount of time was added to the heating cycle to ensure complete disinfection.

**Figure 4.**
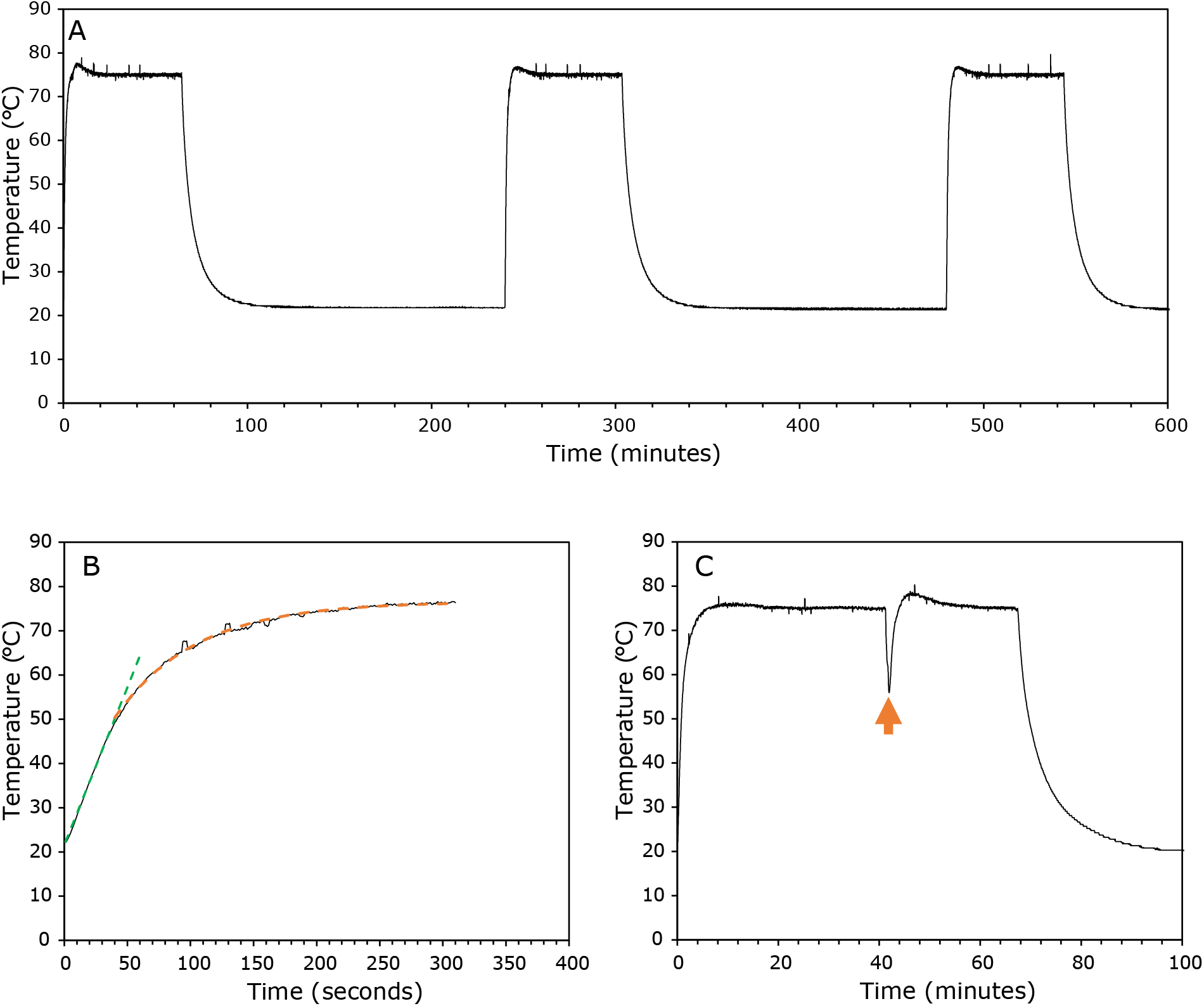
TPH temperature control. A: Three cycles of the TPH heating the tailpiece from room temperature to its target temperature of 75 °C for one hour, followed by 3 hours without heating. This record has been filtered to 0.08 Hz from an original sampling frequency of 1 Hz to reduce noise from the temperature sensor. B: Detail view of the first heating cycle in panel A. The green dashed line is a best fit linear equation to the initial, linear rise in temperature (0.7 °C/sec slope), while the orange dashed line is a fit of temp = *A*+ *B*(1 − *e*^−t /τ^), yielding τ = 64 sec for the exponential phase. C: Response of the TPH to a temperature decrease caused by water flow at the time marked by the orange arrow.

During a typical heating cycle, temperatures of the TPH and the exposed portions of the tailpiece and connected plumbing are well below those that would cause immediate scalds. The hottest external locations during a heating cycle are on the tailpiece directly above and below the TPH, which are heated via conduction from the length of the tailpiece inside the TPH. After forty-five minutes of heating, the temperature of the tailpiece at the threads reaches about 60° C, and the bottom of the tailpiece near the p-trap reaches approximately 54° C. The most accessible point to patients and staff is the upper surface of the drain which remains no hotter than ∼40° C when installed on a basin. Only prolonged contact with these exposed regions would cause first-degree burns (18).

To simulate an event in which the heat tape receives the maximum current from its power supply for an extended duration, a mock device without hardware fail safes was supplied with 12 volts and 2.5 amps. Two trials were conducted to measure the maximum temperatures reached on the inside wall of the tailpiece (128.7 °C), and between the Kapton polyimide tape and the silicone foam insulation (199.6 °C). Despite these high temperatures, the protective outer case remained cool to the touch and experienced only minor melting of its innermost rim (which directly contacts the tailpiece). These results indicate that even in the event of a worst-case device malfunction, the outer case will still be safe to touch and remain intact. Moreover, we confirmed that the thermal cutoff fuse permanently interrupts heating in this unlikely scenario.

#### Determination of effective operational temperature (65°C vs 75°C)

The first set of experiments was conducted with the prototype of the heater using GFP*-E. coli* and two temperature settings of the TPH 65°C and 75°C. In the control sink (without heater), GFP-*E. coli* was able to grow upwards from the p-trap, along the tailpiece and colonize the drain in 7 days – a rate of approximately 25 mm per day. GFP-*E. coli* was consistently detected in the tailpiece and the drain of the control sink (without heater) from day-7 to day-28 (Fig 5). TPH set to 75°C was effective in preventing upward growth and maintaining the sink drain free from GFP-*E. coli* colonization (Fig. 5). After each of the three independent trials, GFP*-E. coli* were consistently present below the tailpiece heater in the all sinks, however, GFP*-E. coli* were not detected in the tailpiece above the TPH set to 75°C or at the drain to which the tailpiece was connected. In contrast, a TPH set to 65°C was not found to be effective in preventing upward growth. GFP-*E. coli* was detected in the tailpiece above the TPH set to 65°C and drain (Fig. 5-Days 14, 21 and 28). These findings show that a heating cycle of 1h at 75°C followed by 3h non-heating period was effective.

**Figure 5.**
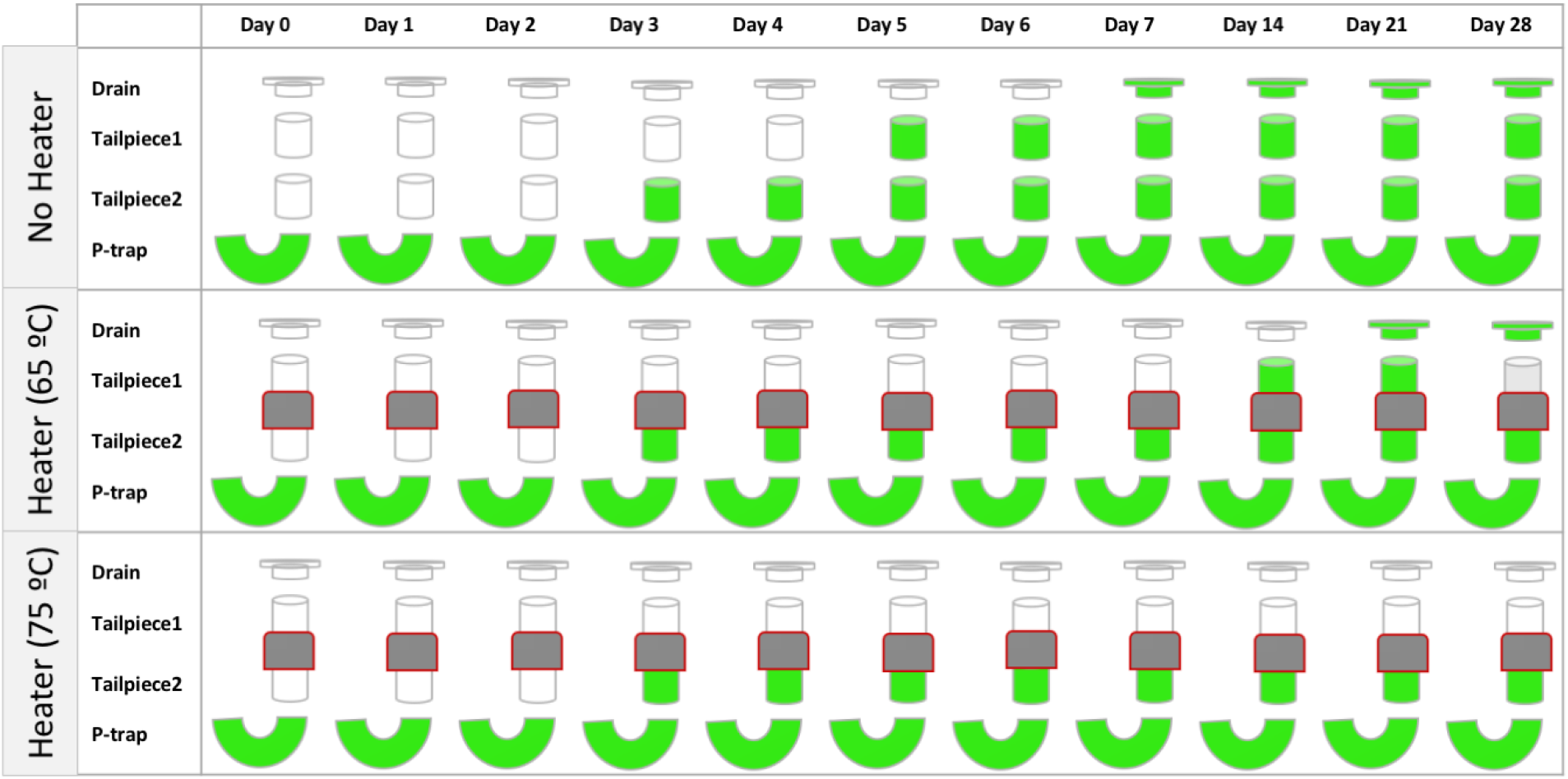
Graphical representation of colonization over time at each of the sampling locations - drain, tailpiece 1(1 inch above heater), tailpiece 2(1inch below heater) and p-trap sections of the sink plumbing. Top most panel without heater and bottom panels with heater set to 65°C and 75°C. Green-filled sections of the plumbing represent detection of GFP-expressing *E*.*coli*.

#### Efficacy against KPCE

With the temperature and cycling time evaluated, a second prototype of the TPH was subsequently challenged with KPCE. In control sinks (without heater), upward growth of KPCE from p-trap to the drain was observed by week 5 (Fig. 6-Sink1 and Sink3). In comparison, KPCE were not detected at the drain or tailpiece above the TPH in sinks with the TPH installed (Fig. 6-Sink2 and Sink4). The TPH were able to keep the sink drains free of KPCE for the 13 continuous weeks of the duration of the test. KPCE were able to persist in the p-trap of all the sinks with or without TPH.

**Figure 6.**
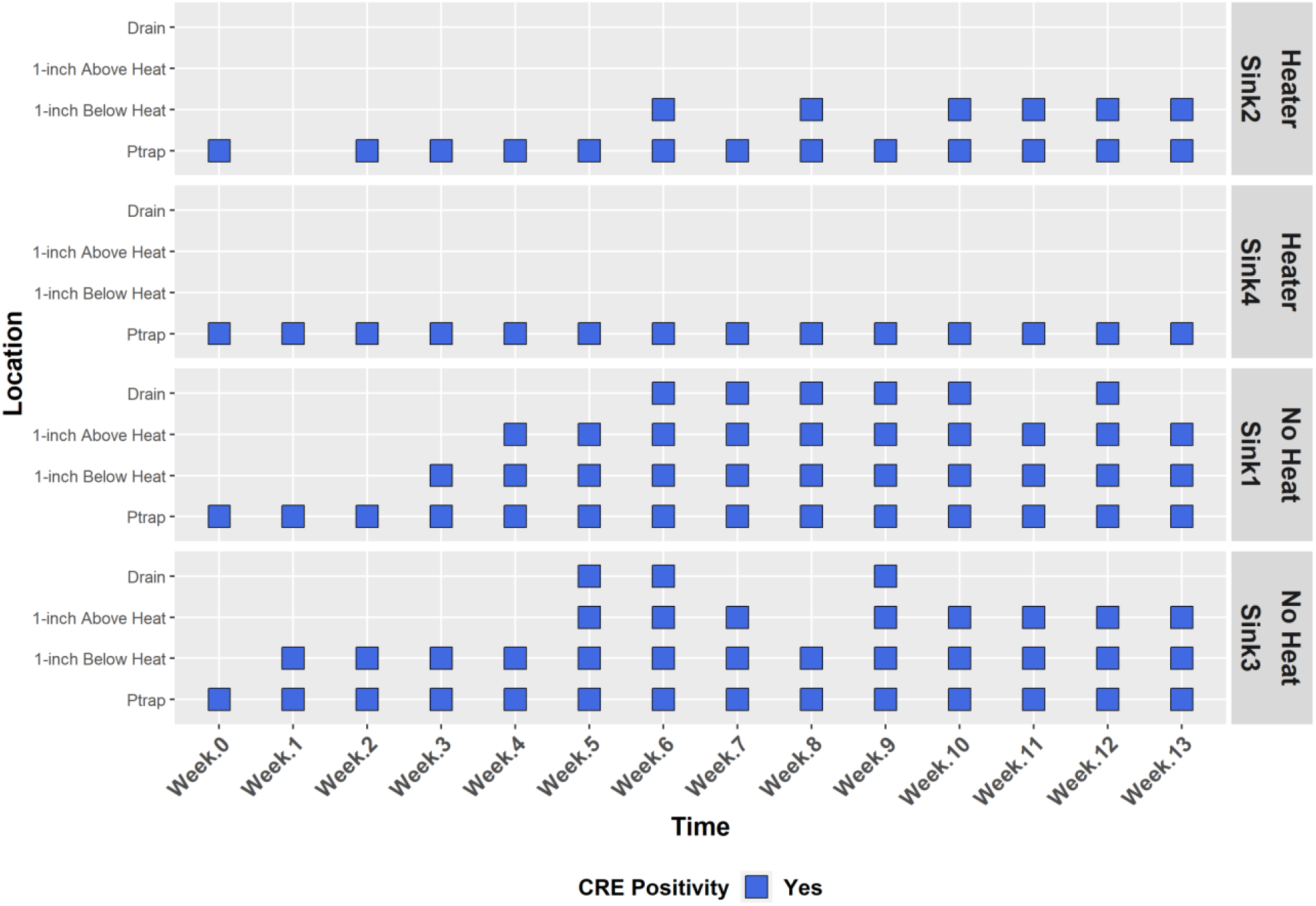
Efficacy of the tailpiece heater in preventing upward growth of KPCE in sinks. Blue cells represent at least 1 CFU of KPCE detected at the corresponding location. Only P-traps in all the sinks were inoculated with KPCE species, new and contamination-free drains and tailpieces were installed at the start (Week 0).

#### Efficacy against KPCE seeding from above

To assess the seeding from above which is likely to happen in a hospital environment, KPCE was seeding from above via the sink drain. Compared to the control sink (without TPH), KPCE was not detected in the sinks with TPH at the level of the drain and the tailpiece beyond a day following inoculation from above (Fig. 7). The TPH was able to maintain the drain and the tailpiece free of KPCE for 34 days tested in this testing. Further, KPCE were not found to establish in the p-trap of the sink with the heater installed. Other bacteria were occasionally detected at the drain and the p-trap in the TPH test sink. In contrast, KPCE were almost consistently detected in the drain, tailpiece and the p-trap of the control sink (without TPH).

**Figure 7.**
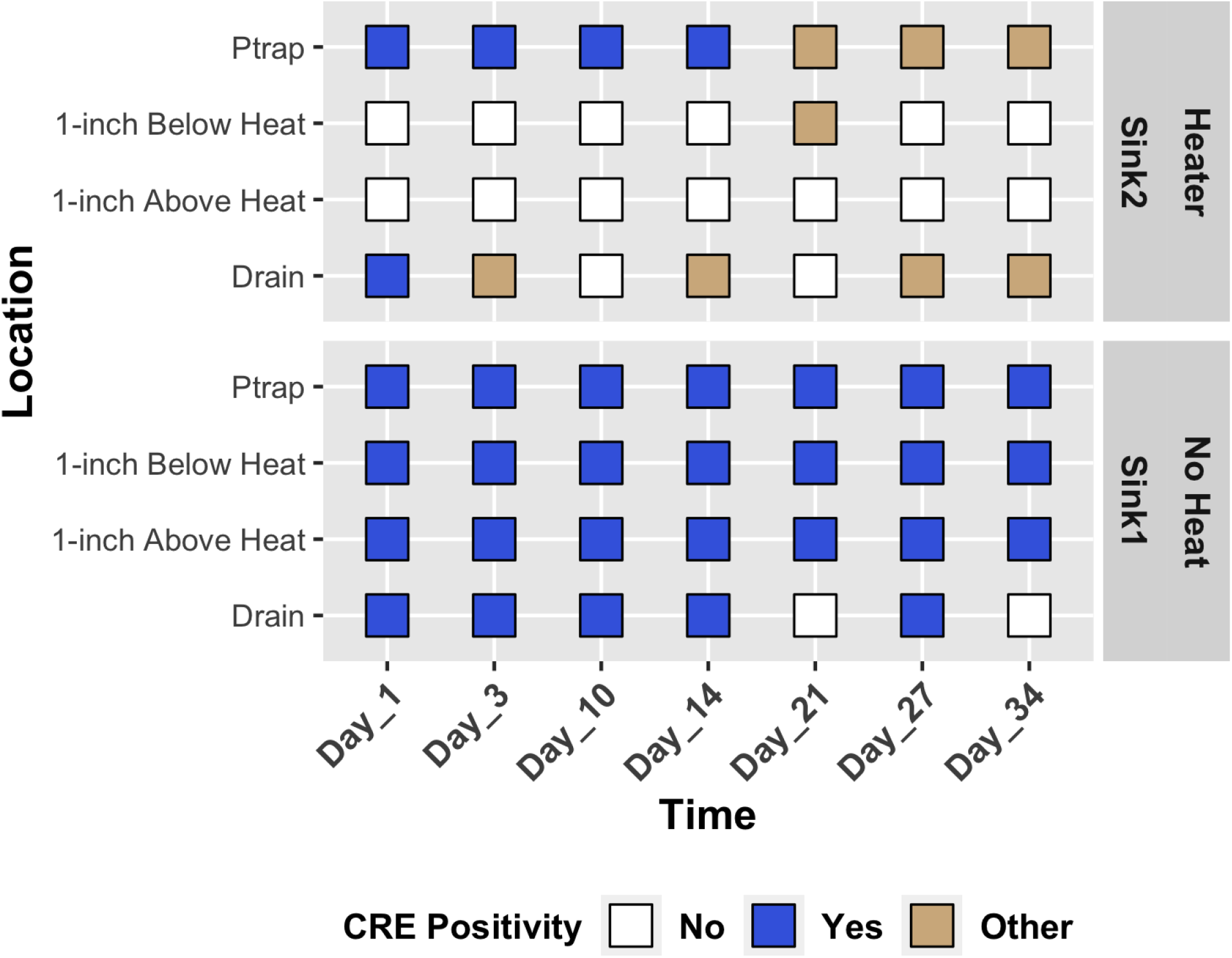
Efficacy of Tailpiece heater in preventing growth of KPCE in sink plumbing following seeding from above. Blue cells represent at least 1 CFU of KPCE detected; White cells represent no growth detected and yellow cells represent at least 1 CFU of *Pseudomonas* or *Stenotrophomonas* species detected. At the start (Day_0) all sink drains were inoculated with KPCE.

## DISCUSSION

The use of heat to prevent upward growth of biofilms from the P-trap to the sink drain and basin is not entirely new. Indeed, others have disclosed the operational principle through patents (19, 20) [US Patents-US4192988A, US3985994A]. To our knowledge, though, the efficacy of these approaches in limiting microbial growth has not previously been tested. Neither has cyclic heating to a minimum sterilization temperature been attempted. The TPH described here heats a 7 cm section of the tailpiece of a sink to 75°C for one hour, followed by 3 hours of non-heating allowing for return to room temperature. Cyclic heating is expected to increase time to device failure, and reduces energy consumption. Total energy cost per sink, per year is estimated to be less than $3. Safety measures are also incorporated and described in the device such as a thermal cutoff to prevent runaway heating as well as insulation.

The effectiveness of cyclical heating in preventing drain colonization was tested using multiple challenge organisms and scenarios. We found that cyclical tailpiece heating to 75°C is effective in preventing colonization of the sink drain by GFP-*E*.*coli* as well as KPCE whether as a result of seeding from above or retrograde from a colonized p-trap. The sink drain itself reaches only 40°C when the tailpiece inside the TPH is at 75°C. Heating the tailpiece is expected to prevent upward growth of organisms from the p-trap, but it is reasonable to question how the non-sterilizing temperature of 40°C could be effective at preventing colonization of the drain when organisms are introduced from above. Anecdotally we find that the drain is kept relatively dry by indirect hearing by the TPH, perhaps presenting an inhospitable environment for biofilm growth. Besides the mechanism of killing by heat the TPH, we speculate that the dissipated heat from the TPH allows the drying of the preexisting biofilm and formation of new biofilm even up to the level of the sink drain.

Our data further show that colonization of the attached p-trap by seeding from above may be reduced by heating the tailpiece. In contrast, seeding of sinks by patients was found to be an issue in a hospital unit where 5% of new drains were found positive when sampled following p-trap heater installation (11). The introduction of new organisms into the sink basin, as would happen when health care workers wash their hands after caring for infected or colonized patients, was the presumed cause.

While interventions such as sink plumbing replacement or chemical agents (bleach, acetic acid etc.) may temporarily reduce sink colonization, these are not practical long-term solutions to the actual problem. Likewise, the above-mentioned p-trap heating device proved not to be a long-term solution. Liquid in the p-trap was subjected to heat and ultrasonic oscillation (11, 13), and after two years there was build-up of scale inside these devices, leaks and device heating failures including overheating which resulted in the removal of all 23 devices from the clinical setting(data not shown). Further, they were found ineffective in preventing drain colonization and therefore potentially ineffective at preventing transmissions from sinks to the surrounding environment and patients (11, 21).

In contrast, the TPH device tested in this study differs in both design and principle from the self-disinfecting p-trap heating device. The TPH device does not directly heat the liquid sitting in the p-trap of the sink, but rather heats the surface of the tailpiece and intends to provide a thermal barrier against biofilms spreading in a retrograde manner along the plumbing surfaces. As such, the TPH does not seek to sterilize incoming contaminated waste from the sink basin. Furthermore, the TPH is relatively small, minimizes power consumption, and doesn’t interfere in any way with routine sink disinfection and hygiene practices.

## ACKNOWLEDGEMENTS

The authors wish to acknowledge a team of students in the University of Virginia class “BME Design and Discovery” who first investigated the approach to heating a tailpiece in 2015 as part of their curricular requirements, and did so with aplomb – Guranchal Hayer, Tom Zanger, Alex Lord, Christopher Winans, Arleigh Grecco, Hans Prakash, and Jordan Hubbell.

## REFERENCES

1. Gordon A, Mathers A, Cheong E, Gottlieb T, Kotay S, Walker A, Peto T, Crook D, Stoesser N. 2017. The Hospital Water Environment as a Reservoir for Carbapenem-Resistant Organisms Causing Hospital-Acquired Infections-A Systematic Review of the Literature. Clinical Infectious Diseases 64:1435–1444.

2. Parkes LO, Hota SS. 2018. Sink-Related Outbreaks and Mitigation Strategies in Healthcare Facilities. Curr Infect Dis Rep 20:42.

3. Carling PC. 2018. Wastewater drains: epidemiology and interventions in 23 carbapenem-resistant organism outbreaks. Infect Control Hosp Epidemiol 39:972–979.

4. French G, Shannon K, Simmons N. 1996. Hospital outbreak of Klebsiella pneumoniae resistant to broad-spectrum cephalosporins and beta-lactam-beta-lactamase inhibitor combinations by hyperproduction of SHV-5 beta-lactamase. J Clin Microbiol 34:358–363.

5. Hota S, Hirji Z, Stockton K, Lemieux C, Dedier H, Wolfaardt G, Gardam M. 2009. Outbreak of Multidrug-Resistant Pseudomonas aeruginosa Colonization and Infection Secondary to Imperfect Intensive Care Unit Room Design. Infect Control Hosp Epidemiol 30:25–33.

6. Gbaguidi-Haore H, Varin A, Cholley P, Thouverez M, Hocquet D, Bertrand X. 2018. A Bundle of Measures to Control an Outbreak of Pseudomonas aeruginosa Associated With P-Trap Contamination. Infect Control Hosp Epidemiol 39:164–169.

7. Stjarne Aspelund A, Sjostrom K, Olsson Liljequist B, Morgelin M, Melander E, Pahlman LI. 2016. Acetic acid as a decontamination method for sink drains in a nosocomial outbreak of metallo-beta-lactamase-producing Pseudomonas aeruginosa. J Hosp Infect 94:13–20.

8. Tofteland S, Naseer U, Lislevand J, Sundsfjord A, Samuelsen O. 2013. A Long-Term Low-Frequency Hospital Outbreak of KPC-Producing Klebsiella pneumoniae Involving Intergenus Plasmid Diffusion and a Persisting Environmental Reservoir. Plos One 8.

9. Vergara-Lopez S, Dominguez M, Conejo M, Pascual A, Rodriguez-Bano J. 2013. Wastewater drainage system as an occult reservoir in a protracted clonal outbreak due to metallo-beta-lactamase-producing Klebsiella oxytoca. Clin Microbiol Infect 19:E490–E498.

10. Volling C, Ahangari N, Bartoszko JJ, Coleman BL, Garcia-Jeldes F, Jamal AJ, Johnstone J, Kandel C, Kohler P, Maltezou HC, Maze dit Mieusement L, McKenzie N, Mertz D, Monod A, Saeed S, Shea B, Stuart RL, Thomas S, Uleryk E, McGeer A. 2021. Are Sink Drainage Systems a Reservoir for Hospital-Acquired Gammaproteobacteria Colonization and Infection? A Systematic Review. Open Forum Infectious Diseases 8.

11. Mathers AJ, Vegesana K, German Mesner I, Barry KE, Pannone A, Baumann J, Crook DW, Stoesser N, Kotay S, Carroll J, Sifri CD. 2018. Intensive Care Unit Wastewater Interventions to Prevent Transmission of Multispecies Klebsiella pneumoniae Carbapenemase-Producing Organisms. Clin Infect Dis 67:171–178.

12. Cole K, Talmadge JE. 2019. Mitigation of microbial contamination from waste water and aerosolization by sink design. Journal of Hospital Infection 103:193–199.

13. Fusch C, Pogorzelski D, Main C, Meyer C, el Helou S, Mertz D. 2015. Self-disinfecting sink drains reduce the Pseudomonas aeruginosa bioburden in a neonatal intensive care unit. Acta Paediatrica 104:e344–e349.

14. Kotay S, Chai W, Guilford W, Barry K, Mathers A. 2017. Spread from the Sink to the Patient: In Situ Study Using Green Fluorescent Protein (GFP)-Expressing Escherichia coli To Model Bacterial Dispersion from Hand-Washing Sink-Trap Reservoirs. Applied and Environmental Microbiology 83.

15. Kotay SM, Donlan RM, Ganim C, Barry K, Christensen BE, Mathers AJ. 2019. Droplet-Rather than Aerosol-Mediated Dispersion Is the Primary Mechanism of Bacterial Transmission from Contaminated Hand-Washing Sink Traps. Applied and Environmental Microbiology 85:e01997–18.

16. Aranega-Bou P, Ellaby N, Ellington MJ, Moore G. 2021. Migration of Escherichia coli and Klebsiella pneumoniae Carbapenemase (KPC)-Producing Enterobacter cloacae through Wastewater Pipework and Establishment in Hospital Sink Waste Traps in a Laboratory Model System. Microorganisms 9.

17. Kotay SM, Parikh HI, Barry K, Gweon HS, Guilford W, Carroll J, Mathers AJ. 2020. Nutrients influence the dynamics of Klebsiella pneumoniae carbapenemase producing enterobacterales in transplanted hospital sinks. Water Research 176:115707.

18. Moritz AR, Henriques FC. 1947. Studies of Thermal Injury: II. The Relative Importance of Time and Surface Temperature in the Causation of Cutaneous Burns. Am J Pathol 23:695–720.

19. Eloranta K, Lindroos R, Surakka J. 1976. Drain pipe sterilization. Finland.

20. Pederson Jr. PD, Ufford KA, Nelson LV. 1980. Electrically heated thermal microbial drain barrier. US.

21. Regev-Yochay G, Smollan G, Tal I, Pinas Zade N, Haviv Y, Nudelman V, Gal-Mor O, Jaber H, Zimlichman E, Keller N, Rahav G. 2018. Sink traps as the source of transmission of OXA-48-producing Serratia marcescens in an intensive care unit. Infect Control Hosp Epidemiol 39:1307–1315.

